# Pan-embryo cell dynamics of germlayer formation in zebrafish

**DOI:** 10.1101/173583

**Authors:** Gopi Shah, Konstantin Thierbach, Benjamin Schmid, Anna Reade, Ingo Roeder, Nico Scherf, Jan Huisken

**Affiliations:** Max Planck Institute of Molecular Cell Biology and Genetics, Pfotenhauerstr. 108, 01307 Dresden, Germany; Cancer Research UK Cambridge Institute, University of Cambridge, Robinson Way, CB20RE Cambridge, UK; Institute for Medical Informatics and Biometry, Medical School, TU Dresden, Fetscherstr. 74, 01307 Dresden, Germany; Optical Imaging Centre Erlangen, Friedrich-Alexander-University of Erlangen-Nuremberg, 91054 Erlangen, Germany; Cardiovascular Research Institute, University of California, San Francisco, CA 94158-9001, USA; Department of Biochemistry and Biophysics, University of California, San Francisco, CA 94158-2517, USA; Morgridge Institute for Research, Madison, Wisconsin 53715, United States of America

## Abstract

Cell movements are coordinated across spatio-temporal scales to achieve precise positioning of organs during vertebrate gastrulation. In zebrafish, mechanisms governing such morphogenetic movements have so far only been studied within a local region or a single germlayer. Here, we present pan-embryo analyses of fate specification and dynamics of all three germlayers simultaneously within a gastrulating embryo, showing that cell movement characteristics are predominantly determined by its position within the embryo, independent of its germlayer identity. The spatially confined fate specification establishes a distinct distribution of cells in each germlayer during early gastrulation. The differences in the initial distribution are subsequently amplified by a unique global movement, which organizes the organ precursors along the embryonic body axis, giving rise to the blueprint of organ formation.

---

An embryo is a product of molecules specifying cells, cells forming tissues and tissues morphing into organs. To understand how information flows across these scales to form and position organs in a developing embryo, it is crucial to have a pan-embryo view of cell and gene expression dynamics. In zebrafish, through the process of gastrulation, morphogenetic movements dynamically organize a mass of undifferentiated cells into distinct germlayers, *ectoderm, mesoderm* and *endoderm*, laying out the primary body plan: cells spread over a spherical yolk during *epiboly*, a subset of which internalize through *involution*, followed by dorsal *convergence* and anterior-posterior (A-P) *extension* of all three germlayers^1^ (fig. S1). Despite an appreciation of their spatio-temporal overlap and remarkable coordination to accomplish gastrulation, each movement and germlayer have so far only been studied separately within a local region or a temporal window of interest^2-5^. Hence regardless of the advances in *in toto* imaging of live embryos^6^, we still fail to comprehend how such a multi-scale process operates as a whole^7^.

Several transgenics and mutants^8,9^ have been used to study the formation of germlayers on the basis of cell behavior, signaling and physical properties^1,10-12^. There is strong evidence that specific cell properties (differential adhesion and surface tension) ^10,11^ and behaviors (random vs. directed cell movement, division and intercalation) ^4,13-16^ drive the formation of individual germlayers. However given the invasive nature of some of the approaches used, visualizing the interplay of all germlayers within an intact developing embryo has remained out of reach. Therefore, it can only be speculated how dynamics within individual germlayers are coordinated across tissues to drive embryo formation. We addressed this by establishing a workflow for non-invasive simultaneous imaging of all three germlayers. Through analyses of embryo-wide gene expression domains and cell movements, we show that a spatially distinct distribution of cells is established within each germlayer during early gastrulation, which is then amplified by a global morphogenetic movement to position organ progenitors along the embryonic body axis.

Early blastoderm cells are equivalent in their genetic composition and expression, so it has been difficult to generate germlayer-specific transgenic labels. To identify all three germlayers, we co-expressed three fluorescent reporters in single zebrafish embryos: *Tg(sox17:H2B*-*tBFP)* (endodermal marker), *Tg(mezzo:eGFP)^17^* (pan-mesendodermal marker) and *Tg(h2afva:h2afva*-*mCherry)*^18^ (ubiquitous nuclear marker). A custom 4-lens Selective Plane Illumination Microscopy (SPIM) setup^19^ was used to perform *in toto* imaging of the triply labeled embryos with high acquisition speed (~30s per time-point). The amount of data was reduced by acquiring only a 300 *μ*m thick spherical shell around the embryo surface (fig. S2). The raw data from all three channels were then computationally separated into three germlayers, depending on the presence of specific reporters-prospective ectoderm (includes the EVL and YSL cells) expressed only *histone*, mesendoderm expressed *histone* and *mezzo*, while endoderm expressed all three reporters (fig. S3; movie S1; methods).

Visualizing the spatio-temporal dynamics of *mezzo* expression, one of the first nodal-activated transcription factors expressed in mesendoderm^20^, showed that mesendoderm specification is a gradual process beginning at 4.5 hpf at the dorsal lip and spreading laterally to cover the entire germ ring (Fig. 1A). As mesendoderm cells were specified, the remaining blastoderm cells were used as an indirect readout of ectoderm specification, termed here as prospective ectoderm (p. ectoderm; these cells can still acquire mesendodermal fate) (Fig. 1B,F). At 6.5 hpf, the dorsal forerunner cells (DFC), the only non-invaginating cells of endodermal origin expressing *sox17*, were specified, and the rest of the endoderm formed from the internalized mesendoderm pool (Fig. 1C,F). We detected these cells and quantified the relative abundance of cells in each germlayer from 4-12 hpf, which provided a comprehensive, quantitative depiction of cell fate specification. Relative cell counts of the three germlayers reached a plateau by 9 hpf, marking the completion of specification (Fig. 1E). All germlayers thereafter engaged in convergence-extension movements. Interestingly, p. ectoderm cells moved to the anterior of the embryo while converging towards dorsal to form the brain, notochord and optic cups (Fig. 1B,D). Mesendoderm cells converged towards mid-dorsal as well as anterior and posterior parts, with mesoderm giving rise to somites (Fig. 1A,D) and endoderm forming the lining of gut along with the assembly of Kupffer’s vesicle (Fig. 1C,D). Thus, imaging embryo-wide expression patterns of *histone*, *mezzo* and *sox17* provided simultaneous visualization of specification and dynamics of all germlayers within the developing embryo (movie S1, S2).

**Figure 1.**
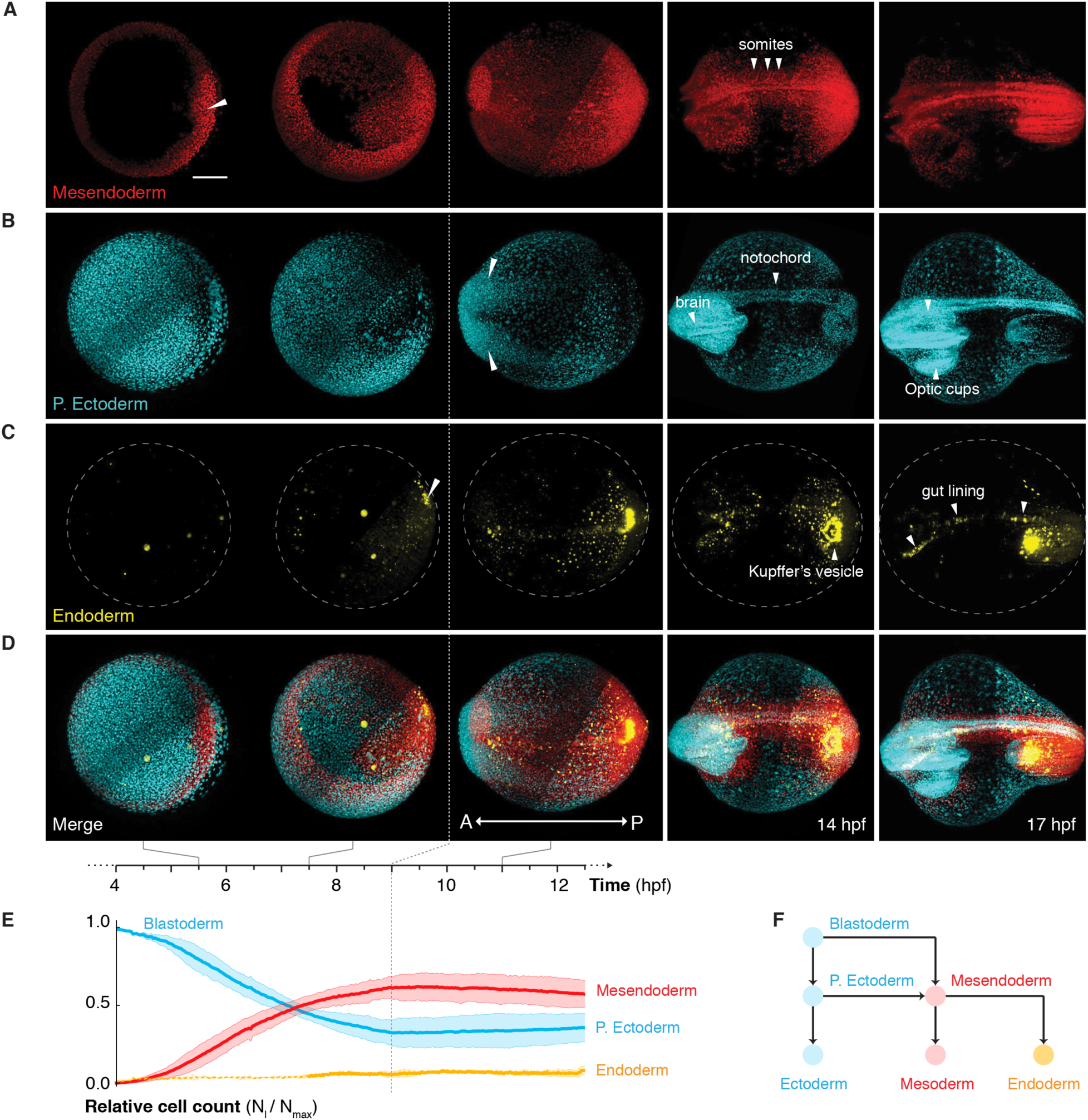
*In toto* imaging of germlayer specification and dynamics. (**A-C**) Fate specification and dynamics of mesendoderm (red), p. ectoderm (cyan) and endoderm (yellow) cells spanning 4-17 hpf. (**A**) Mesendoderm specification begins at the dorsal lip (white triangle), spreads around the germ ring and converges towards the dorsal midline, forming the somites (white triangles). (**B**) P. ectoderm cells converge towards the anterior (white triangles) leading to formation of brain, notochord and optic cups. (**C**) Endoderm specification begins at the dorsal shield (white triangle) with the DFCs, followed by rest of the endoderm, which forms the gut lining and Kupffer’s vesicle upon dorsal convergence. (**D**) Merge of all three germlayers. Scalebar: 200 μm. A: anterior, P: posterior (**E**) Mean relative cell numbers (n=3) for p. ectoderm (blue), mesendoderm (red) and endoderm (yellow). Bands correspond to 1.96**\***standard error. Dashed part of the yellow line indicates the period before endoderm specification. (**F**) Schematic explaining germlayer specification showing blastoderm differentiating into mesendoderm and p. ectoderm. Mesendoderm further differentiates into mesoderm and endoderm.

We hypothesized that such complex organization of cells during gastrulation is accomplished in two conceptual phases: (i) radial organization to stratify blastoderm cells into germlayers and (ii) axial organization to position various organ progenitors at specific locations along the body axis. To this end, we segmented and tracked the cells in each germlayer to investigate the radial and axial organization of cells (movies S4, S5). Cell trajectories depict the cellular flows of individual germlayers and their mutual coordination throughout the process (movies S5, S6).

During early gastrulation (4-7 hpf), a subpopulation of *mezzo* expressing cells involuted at 5.5 hpf forming a multi-layered mesendoderm. Subsequently, they moved towards the animal pole, sliding along the outer ectodermal cells undergoing epiboly^2^ (Fig. 2A,B). All three germlayers continued their epiboly movement towards the vegetal pole to spread over the yolk (Fig. 2B). Though endoderm cells were specified around 6.5 hpf, they retained their salt and pepper distribution with mesoderm cells as previously reported^21^, and separated from the mesoderm at the end of epiboly (Fig. 2B-D; fig. S4; movie S3). Through this complex execution of epiboly and internalization of cells alongside fate specification, the desired radial stratification and thinning of layers was accomplished by ~9 hpf, as shown by the radial position of germlayers normalized to the mesendoderm position at each time-point (Fig. 2E).

**Figure 2.**
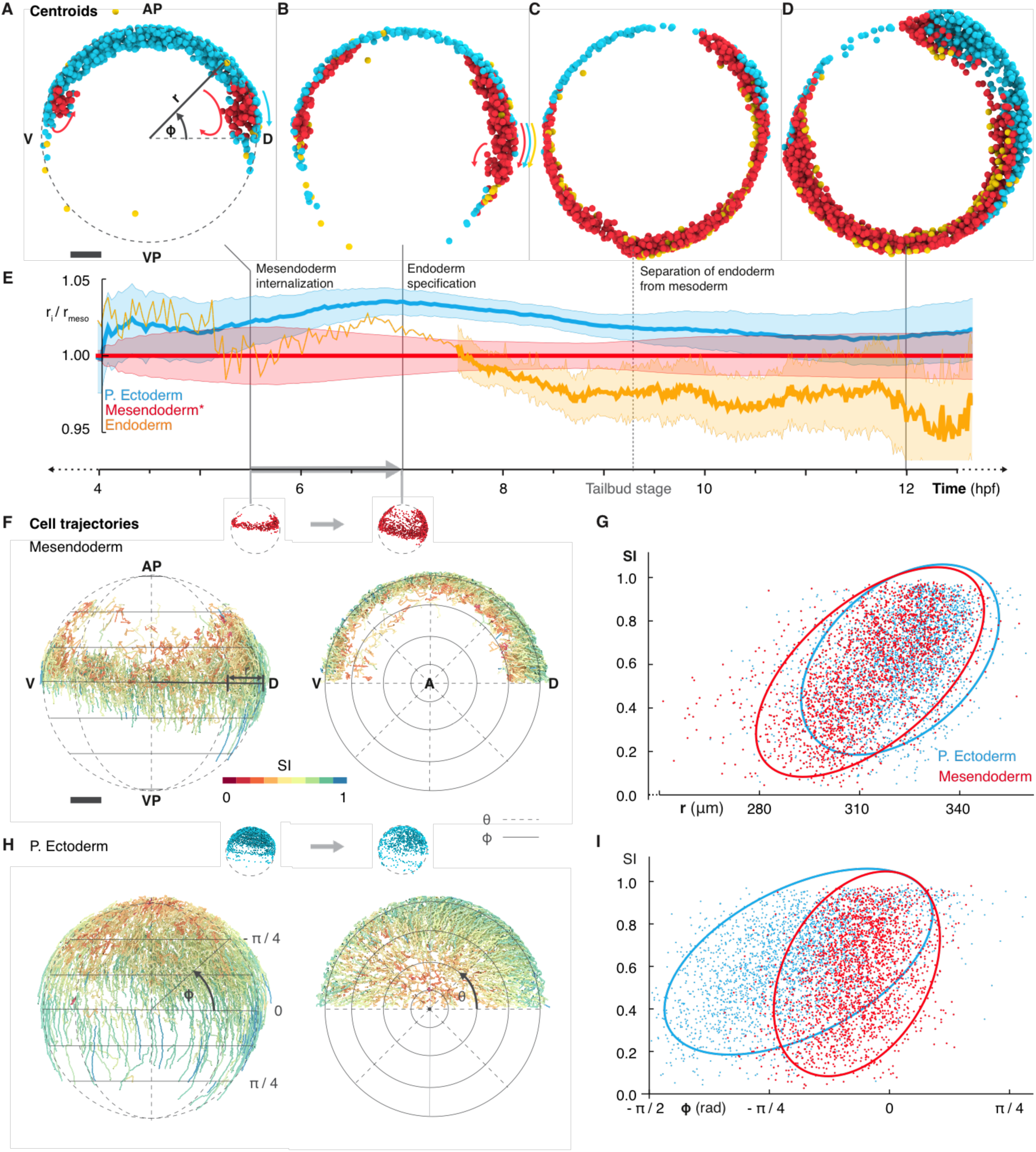
Position dependent organization of cell movement during early gastrulation. (**A-D**) Lateral views of the embryo at 4, 6.5, 9 and 11.5 hpf showing rendered object centroids located within a plane through the body axis (+- 78 μm). Colors indicate the corresponding germlayer: p. ectoderm (blue) mesendoderm (red) and endoderm (yellow). Colored arrows in (**A**) indicate epiboly of p. ectoderm (blue) and internalization of mesendoderm (red), in (**B**) indicate epiboly movement of all three germlayers. AP: animal pole, VP: vegetal pole, D: dorsal, V: ventral (**E**) Line plot showing mean radial position of all germlayers (normalized with respect to average radius of the mesendoderm) for a single embryo. Bands indicate mean +/- 0.3 standard deviation, the scaling was introduced to reduce overlap between germlayers and to visually highlight the thinning of layers. Dashed yellow line indicates the period before endoderm specification. (**F**) Cell trajectories for mesendodermal cells during early gastrulation (4.5 - 7 hpf; indicated by gray arrow in (**E**)) shown in lateral (left) and animal-pole view (right). Color code indicates the straightness indices (SI) of trajectories. (**G**) Scatterplot of SI vs. radial position r (computed at the mid-point) for each trajectory (4.5 - 7 hpf) of mesendoderm (red), p. ectoderm (blue) and the respective 90% prediction ellipsoids. (**H**) Cell trajectories for p. ectoderm, same views and color code as in (**D**). (**I**) Scatterplot of SI vs. position ϕ along the longitude (computed at the mid-point) for each trajectory (4.5 - 7 hpf) of mesendoderm (red), p. ectoderm (blue) and the respective 90% prediction ellipsoids.

Several attempts have been made to find an order in this complex process, assuming germlayer specific cell behaviors: the endoderm has been shown to perform random walk after involution followed by directed dorsal-ward movement^4^, the ectoderm is thought to perform cell intercalation and collective cell movement^1,14,15^, whereas the mesoderm has been shown to exhibit a variety of behaviors along the dorso-ventral axis to pattern its motion^12-14,16^. To assess if the characteristic of cell movement (random vs. directed) was indeed germlayer specific, we measured how straight cells in each germlayer moved and correlated the straightness indices (SI) of cell trajectories to each of *r* (radius), *θ* (latitude) and *ϕ* (longitude) coordinates (methods). We found a striking pattern: both mesendoderm and p. ectoderm cell tracks closer to the embryo surface (large *r*) exhibited a higher SI as compared to those at deeper locations (small *r*), revealing a radial organization (Fig. 2F-H). Likewise, p. ectoderm cell tracks close to the animal pole (*ϕ* < -*π*/4) showed a lower SI than cells closer to the margin (*π*/4 < *ϕ* < -*π*/4), whereas no obvious pattern was observed along *θ* (Fig. 2H,I; fig. S5), indicating that the ectoderm cell movement is also organized along the animal-vegetal axis of the embryo. This global perspective of single cell trajectories uncovered that irrespective of their germlayer identity, the straightness of cell movement strongly correlates with the cell’s position within the embryo (Fig. 2G,I), implying an emergent position dependent, rather than a germlayer specific, pattern in cell movement that drives local and global cell reorganization during early gastrulation.

While the movement characteristics of cells appeared to be independent of germlayer identity, we found a unique distribution of cells in each germlayer at the end of early gastrulation, depicted by cell density rendering showing medio-posteriorly located mesendoderm and anteriorly located p. ectoderm cells (Fig. 3A). This suggested that the pattern of initial fate specification is a crucial event in establishing the distribution of cells within germlayers. However, it still remained to be understood if germlayer dynamics are regulated individually or a in a global fashion. To investigate this, we decomposed trajectories^22^ of every cell into an epiboly component (movement from animal to vegetal pole) and a convergence component (movement from ventral to dorsal side of the embryo) throughout development. In a spherical coordinate system of a suitably oriented embryo, convergence corresponds to motion along the parallels, whereas epiboly and extension correspond to directed motion along the meridians before and after completion of epiboly, respectively (Fig. 3E’,E”; fig. S6). The radial movement of cells, which does not contribute directly to either of the movement components, was ignored in this analysis. Our results show that while the p. ectoderm undergoes a peak of epiboly followed by a peak of convergence in its movement, the mesendoderm has constant epiboly and convergence components throughout the process (Fig. 3B,C; fig. S7). Therefore we hypothesized that the movements of ectoderm and mesendoderm cells are decoupled during late gastrulation as formerly postulated^1,8,23^ and that each germlayer might possess a characteristic motion pattern as demonstrated for the endoderm in our previous study^19^.

**Figure 3.**
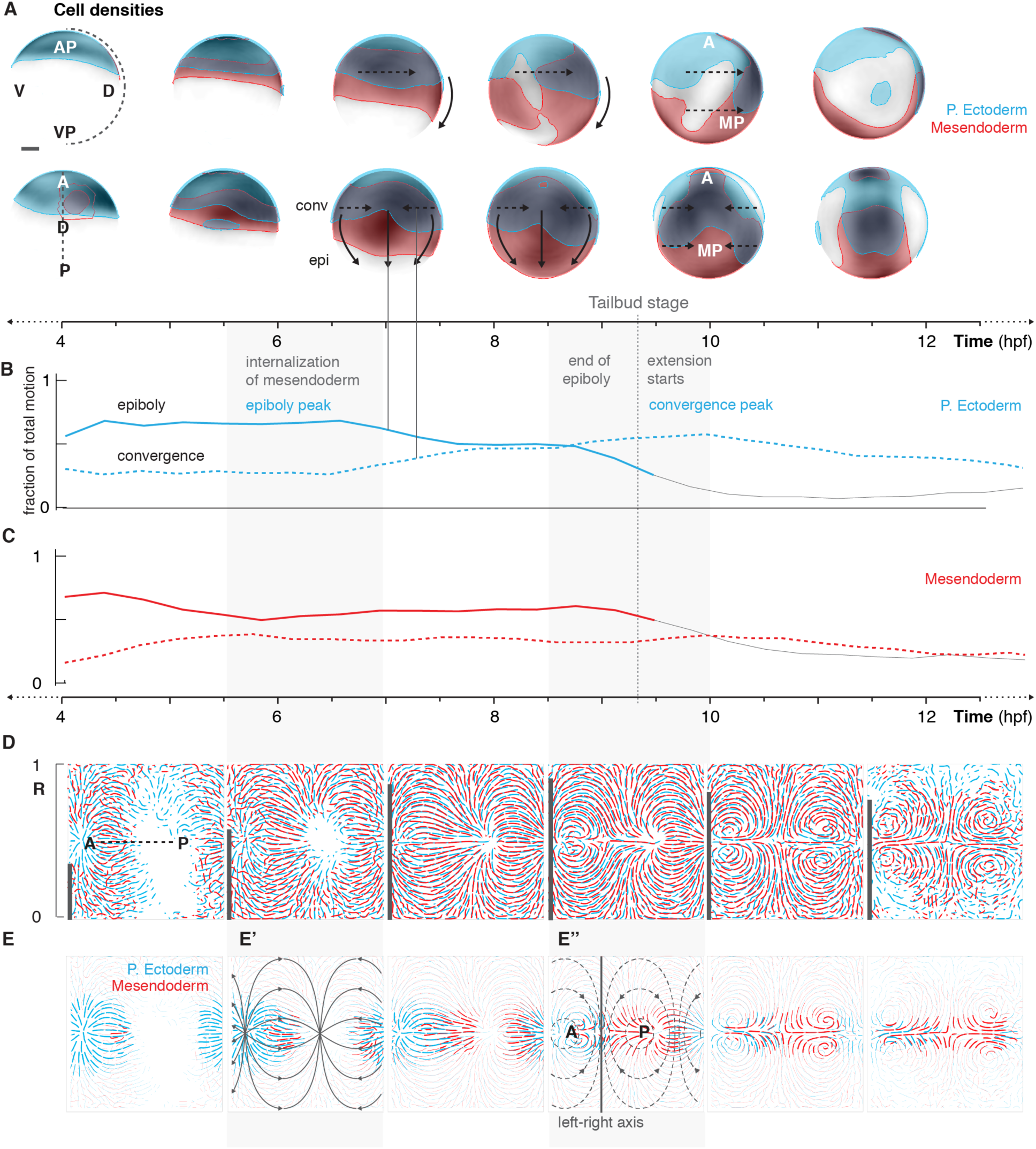
Distinct motion patterns of germlayers during late gastrulation. (**A**) Cell densities shown as grayscale (white-low to black-high) on spherical representation of embryo. Regions of high densities (>0.33 normalized density) are color coded, showing the extent of p. ectoderm (blue) and mesendoderm (red) at various time points. Data shown in lateral (top) and dorsal (bottom) views. Arrows indicate expected direction of epiboly movement (solid) and convergence movement (dashed). AP: animal pole, VP: vegetal pole, MP: medio-posterior (**B-C**) Proportions of epiboly (solid) and convergence (dashed) movement for p. ectoderm (**B**) and mesendoderm (**C**): lines shows mean across n=3 embryos. Gray dashed line indicates end of epiboly (tailbud stage). (**D**) General motion patterns of p. ectoderm and mesendoderm shown as streamlines in Mercator projections, each interval covers ~1.5 hours of development. Dorsal midline indicated by dashed line. Correlation of motion between germlayers is indicated as bars on the left edge of each streamline plot. (**E**) Tissue flow shown as density-weighted streamlines for p. ectoderm and mesendoderm, thickness of streamlines indicates cell density at the respective site. Principal directions for epiboly (solid-gray) and convergence (dashed-gray) movement in Mercator projection are shown as overlay in (**E’**) and (**E”**) respectively. (**E”**) Border separating the ectodermal and mesendodermal flows is indicated by a solid line along the left-right axis.

In order to identify these germlayer specific motion patterns, we merged cell-tracking data from multiple embryos and obtained visualizations of cellular flows^24^ for each germlayer individually (methods). To our surprise, they appeared strikingly similar, with a high correlation between the local flow directions of the mesendoderm and p. ectoderm throughout the second phase of gastrulation (Fig. 3D). Hence, in contrast to our hypothesis, we show that a single unifying motion drives the dynamics of all three germlayers (Fig. 3D; fig. S8). Distinct patterns emerged when the streamlines were scaled to reflect local cell densities of respective germlayers, indicating that the resultant organization of cells in each germlayer is dependent on the cell density distribution (Fig. 3E). Further, we found a separation of flows along the left-right axis between the p. ectoderm and mesendoderm domains, which was strongest around 9 hpf (Fig. 3E”). This boundary explains the prominent convergence of the p. ectoderm towards the anterior and that of mesendoderm towards the medio-posterior region of the embryonic axis, as already apparent in Fig. 1 (Fig. 3E). Taken together, these results show that epiboly, convergence and extension are parts of a major underlying movement that drives large-scale deformations of the embryo. As all germlayers experience the same global movement, the spatio-temporal pattern of cell fate specification and the resulting density distribution of each cell type during early gastrulation are crucial in determining the positioning of organ progenitors at the end of gastrulation.

In summary, we present a pan-embryo visualization of differentiation of blastoderm into three germlayers and their dynamics at cellular resolution. Through assessment of the process at the level of single cells, gene expression domains, entire germlayers and whole embryos simultaneously, we uncovered that during early gastrulation, spatially confined mesendodermal fate specification combined with the position dependent cell movement establishes a distinct initial distribution of cells in each germlayer. The differences in this initial distribution of cell types are amplified during late gastrulation, wherein a unique global movement organizes the organ precursors along the embryonic body axis, independent of their germlayer identity. This entire process gives rise to the blueprint of organ development in zebrafish. While gastrulation has been thought of as a complex process comprising multiple cell behaviors and movements driving different aspects of germlayer organization, our analyses show that it in fact is a phenomenon of whole^7^, governed by regulation of basic parameters such as initial cell positions and their density distribution. Such a mechanism relies on an accurate fate specification in space and time, as displacements in the initial mesendodermal population would get amplified and potentially lead to aberrant development. For example, displacement of endoderm precursor cells at the onset of gastrulation in *cxcr4a* morphants leads to mis-positioning of endoderm-derived organs, despite cell movements being unaffected.^19^

Further, our study illustrates the power of combining genetic tools, high-speed light sheet microscopy and rigorous data analysis to understand how a process like gastrulation is orchestrated across spatio-temporal scales. Being able to visualize and track cell cohorts from their inception to incorporation into organs will benefit developmental and disease-oriented studies^25^. Conventionally, such questions have been addressed through fate mapping experiments using sparse labeling that report spatial position of cells^26^. Here, we demonstrate that our data contain not only the spatial positions but also the temporal information about large-scale cell movements that assemble specific organ precursors to form organs. We believe that such a singular dataset offers a crucial resource for the community to reconstruct the emergence and interplay of specific tissues and organs during early zebrafish development.

## Acknowledgements

We thank Scott Fraser, Vikas Trivedi, Pia Aanstad and Thomas Zerjatke for discussions at various stages of the project. We also thank Rory Power and Gloria Slattum for their comments on the manuscript. Work in the lab of J.H. was supported by the Max Planck Society and European Research Council (CoG SmartMic, 647885). Work in the group of I.R. was funded by the Excellence Initiative by the German Federal and State Governments (Institutional Strategy, measure “support the best”) awarded to I.R.

## Author Contributions

G.S. and J.H. conceived the project. G.S. performed the experiments and post-acquisition image processing. G.S., B.S. and J.H. built the microscope setup. B.S. implemented the microscope control, image acquisition and processing software. K.T. and N.S. implemented the software for data analysis and data visualization. G.S., K.T. and N.S. analyzed the data. K.T. and N.S. visualized the data. A.R. made the *Tg(sox17:H2B*-*tBFP)* transgenic fish line. I.R., N.S. and J.H. supervised the project. G.S., K.T., N.S. and J.H. wrote the manuscript with inputs from all authors.

## Supplementary Materials

Materials and Methods

Fig S1 – S8

References (27 – 33)

Movie S1-S6

